# Rgf1 GEF activity toward Rho1 defines a new actin-dependent signal to determine growth sites independently of microtubules and Tea1

**DOI:** 10.1101/2024.01.10.574961

**Authors:** Patricia Garcia, Ruben Celador, Tomas Edreira, Yolanda Sanchez

## Abstract

Cellular asymmetry begins with the selection of a discrete point on the cell surface that triggers Rho-GTPases activation and localized assembly of the cytoskeleton to establish new growth zones. The cylindrical shape of fission yeast is organized by microtubules that deliver the landmark Tea1–Tea4 complex at the cell tips to define the growth poles. However, only a few *tea1*Δ cells mistaken the direction of growth, indicating that they manage to detect their growth sites. Here we show that Rgf1 (Rho1-GEF) and Tea4 are components of the same complex and that Rgf1 activity toward Rho1 is required for strengthen Tea4 at the cell tips. Moreover, in cells lacking Tea1, selection of the correct growth site depends on Rgf1 and on a correctly polarized actin cytoskeleton, both necessary for Rho1 activation at the pole. We propose an actin-dependent mechanism driven by Rgf1–Rho1 that marks the poles independently of microtubules and the Tea1–Tea4 complex.

## INTRODUCTION

Cell polarity is the primary mechanism for generating cellular asymmetry, which is critical for most cell and tissue functions such as development, cell migration and differentiation in a wide variety of organisms including humans. It typically begins with a signal on the cell surface that triggers a cascade of molecular events that induce the localized assembly of cytoskeletal and signaling networks, which subsequently direct the formation of a new growth area (1,2). Fission yeast has been used over the last decades as a simpler and more accessible model for studying this complex process (3). Its cells have a cylindrical shape that is maintained throughout the entire cell cycle, changing in length but not in diameter. This phenomenon is achieved by restricting growth to the cell poles, a process that is still not well understood. Growth occurring at the cell ends has been mainly studied in the transition from monopolar to bipolar growth, termed New End Take-Off (NETO) (4–7). NETO depends on specific polarity determinants, the kelch-repeat protein Tea1, the SH3 domain-containing protein Tea4 and the DYRK kinase, Pom1 among others (8–11). In the absence of Tea1–Tea4 complex, cells grow monopolarly but maintain their cylindrical shape. However, under certain stresses, the cells often choose the wrong growth site, forming bulged and T-shaped cells (10,12).

Tea1 and Tea4 ride on growing microtubule (MT) plus ends to the cell tips, where they are released as discrete “dots” at the cortex, being Tea4 totally dependent on Tea1 for its location (12–15). At poles, the prenylated protein Mod5 and the ERM (Ezrin-Radixin-Moesin) family protein Tea3 anchor Tea1 to the cell cortex (16,17). Tea1 and Tea4 colocalize at the cell tips to form clusters or nodes, (18) and it is assumed that this association promotes the binding of other polarity factors in large protein complexes that organize polarized growth (4,10,19). One of these proteins is the formin For3, whose association with polarity markers likely brings it into the proximity of activators, stimulating the formation of F-actin cables that will deliver growth cargo to the tip (9).

Establishing polarized growth involves hundreds of proteins; however, a constant from yeast to humans involves local accumulation of active GTP-bound forms of Rho-family GTPases at the cell cortex (20–22). While polarity is mostly associated with the functions of Rac and Cdc42 in mammals or Cdc42 in yeast, other GTPases such as RhoA or Rho1 (yeast) can also play a role in the development of polarity. Active RhoA is found at the leading edge as the edge advances in migrating cells, whereas Cdc42 and Rac1 are activated later (23). In budding and fission yeast, Cdc42 displays the ability to polarize spontaneously (22,24–26). However, the role of Rho1, the other essential GTPase, in polarized growth remains undefined. Depletion of Rho1 in growing cells induces shrinking and death via a kind of “apoptosis” that is accompanied by the disappearance of polymerized actin (27,28). Rho1 activity is regulated by three GEFs, Rgf1, Rgf2, and Rgf3 that catalyze the exchange of GDP for GTP, rendering the GTPase in an active state (28–34). The main activator of Rho1, Rgf1, regulates cell integrity through Rho1 by activating the β-glucan-synthase complex (28) and gene expression via the Pmk1 MAPK cell integrity-signaling pathway (28,35,36). Moreover, Rgf1, like Tea1, Tea4 and other polarity factors, is required for the actin reorganization necessary to switch from monopolar to bipolar growth during NETO (28).

Here we have studied this phenomenon to show that Rgf1 interacts with the cell end marker Tea4 and binds to the plasma membrane (PM) through its PH domain. Both, PM binding and Rho-GEF activity are required for stable accumulation of polarity markers at the cell poles. In addition, we have uncovered a new role for Rgf1 in restricting growth to the poles in the absence of polarity markers. Most *tea1*Δ cells maintain their cylindrical morphology unless subjected to stresses, suggesting that these cells detect the location of its poles by an unknown mechanism. Here we show that this mechanism depends on the actin cytoskeleton and Rho1 activation by Rgf1. Therefore, we propose two parallel pathways to define the growth poles in fission yeast: the canonical one dependent on MTs and Tea1**–**Tea4 and another one dependent on actin and Rgf1–Rho1, both necessary to maintain a straight shape when the other is impaired.

## RESULTS

### Rgf1 is required for proper localization of the Tea1–Tea4 complex

We have previously shown that *rgf1*Δ strain displays defects in bipolar growth, with ∼80% of the cells showing monopolar pattern of growth compared to ∼25% of the wild-type cells (28). This growth defect has been described in mutants affected in polarity factors such as Bud6, Tea1 or For3 (37–39), suggesting that similarly to these proteins, Rgf1 triggers NETO, and thus *rgf1*Δ cells may have problems when choosing the right end of growth. This prompted us to examine the localization of the landmark proteins Tea4 and Tea1 in cells lacking Rgf1. While Tea4-GFP was concentrated at both cell tips in wild-type cells (9,11), in *rgf1*Δ cells the signal detected was visibly diminished. We compared the fluorescence intensity of Tea4-GFP in wild-type and *rgf1*Δ cells in the same preparation, in which we grew *tea4-GFP rgf1^+^ sad1-dsred* (spindle pole body marker) cells and *tea4-GFP rgf1*Δ cells separately, and then mixed and imaged them at the same time (Fig 1A). In *rgf1*Δ cells, the Tea4-GFP signal was dispersed in small dots that spread out at the ends. The average fluorescence of Tea4-GFP dots at the tips of *rgf1*Δ cells was approximately half of that seen at the tips of the wild-type cells. Because the *rgf1*Δ cells grew in a monopolar fashion, we examined whether the Tea4 dots were more prominent at one end or whether they were scattered similarly at both ends. We observed that 61% of the *rgf1*Δ cells accumulated Tea4 at the non-growing end (revealed by calcofluor staining) (Fig 1B). Thus, Rgf1 is more important to localize Tea4 to the growing tip, the one where Rgf1 concentrates in wild-type monopolar cells (Fig S1A).

**Fig 1:**
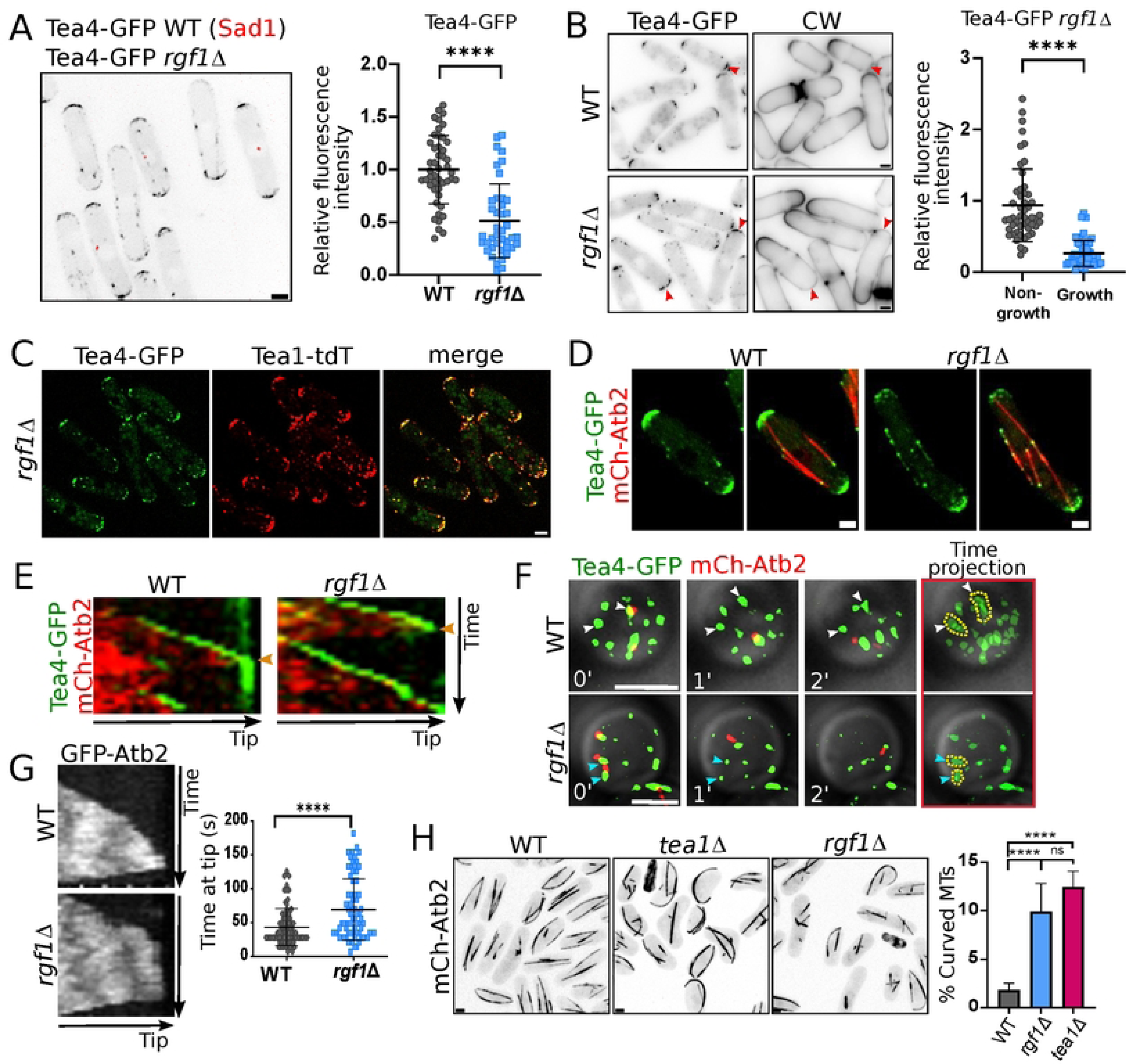
Rgf1 is required for proper localization of the Tea1-Tea4 complex at the cell tip. (A) Cells expressing *tea4*-*GFP rgf1^+^ sad1-dsred* and *tea4-GFP rgf1*Δ were grown in YES liquid medium separately, and then mixed and imaged in the same preparation. The maximum-intensity projection of six deconvolved Z-slides (0.5 µm step-size) is shown. The graphic represents the mean ± standard deviation (SD) of the relative fluorescence intensity of Tea4-GFP measured at the cell tips in the wild-type (WT, n = 47) and *rgf1*Δ (n = 41) cells. WT levels were used for normalization. (B) Calcofluor white (CW, 20 µg/mL) staining and GFP fluorescence in cells expressing *tea4-GFP rgf1^+^* and *tea4-GFP rgf1*Δ. The maximum-intensity projection of six Z-slides (0.5 µm step-size) of Tea4-GFP fluorescence is shown. The red arrowheads indicate non-growing poles. The graphic represents the mean ± SD of the relative fluorescence intensity of Tea4-GFP measured at the tips of the *rgf1*Δ (n = 50) cells. Calcofluor staining, which marks sites of cell growth, was used to differentiate growth from non-growth poles (right). (C) Representative images of Tea4-GFP (green) and Tea1-tdTomato (red) localization in the *rgf1*Δ cells. The maximum-intensity projection of six Z-slides (0.5 µm step-size) is shown. (D) Representative images of the indicated cells expressing *tea4-GFP* (green) and *mCherry-atb2* (red). The maximum-intensity projection of six Z-slides (0.5 µm step-size) is shown. (E) Kymographs of time-lapse fluorescence movies of *tea4-GFP* and *mCherry-atb2* expressed in wild-type or *rgf1*Δ strains. The maximum-intensity projection of seven Z-slides (0.6 µm step-size) of images taken every 15 s was used to draw a line along an MT from the middle of the cell to the tip. The orange arrowheads indicate the moment of MT retraction. (F) Super-Resolution Radial Fluctuations (SRRF) images of Tea4-GFP (green) and mCherry-Atb2 (red) in “head-on” cell tips of the indicated strains. One focal plane image was taken every minute. The time projection of the three images at different time points is shown to follow Tea4 cluster movement (right). Note that in the wild-type strain Tea4 nodes remain stable (white arrowheads) for longer than in the *rgf1*Δ mutant (blue arrowheads). (G) Kymographs of time-lapse fluorescence movies of GFP-Atb2 producing in the WT and *rgf1*Δ cells. The graph shows the mean ± SD of the time during which the MT is touching the pole in both strains (n = 75). (H) Representative images of the indicated cells producing mCherry-Atb2. The maximum-intensity projection of six Z-slides (0.5 µm step-size) is shown. The graph shows the mean ± SD of the percentage of curved MTs found in the indicated strains (n > 500). Statistical significance was calculated using two-tailed unpaired Student’s t test. ****P < 0.0001; ns = non-significant. Scale bar 2 µm.

We also evaluated whether the localization of Tea1, which functions in a complex with Tea4 (9), is affected in the Rgf1 mutant. The wide co-localization between Tea1 and Tea4 in the absence of Rgf1 indicates that both proteins displayed similar localization defects (Fig 1C), and that similarly to wild-type cells, in *rgf1*Δ cells Tea1 and Tea4 were still bound and formed a complex. In *tea4*Δ cells, Tea1 is concentrated at the non-growing cell tip (9). Given that in *rgf1*Δ cells, Tea1 also localized mainly to the non-growing end (Fig 1B and C), our results suggest that the Tea1 mislocalization could be a consequence of Tea4 mislocalization. We noticed that *rgf1*Δ cells showed a greater number of Tea4 discrete dots in the middle zone of the cell (Figs 1A, 1B, and S1B). Examination of Tea4-GFP in wild-type and *rgf1*Δ cells with α–tubulin labeled in red (mCherry-Atb2) showed co-localization of Tea4 cytoplasmic dots with MTs (Fig 1D). Thus, in the *rgf1*Δ cells, a larger free cytoplasmic pool of Tea4, which is not properly sequestered at the poles, could now be available to be redirected to the cell cortex by MTs.

### Rgf1 functions to integrate Tea4 in big clusters at the cell tip

Next, we determined whether Tea4 was accurately delivered to the cell cortex in *rgf1*Δ cells by taking time-lapse images every 15 seconds. We did not observe appreciable differences in the delivery of Tea4 to the cell cortex between *rgf1*^+^ and *rgf1*Δ cells. However, the Tea4-GFP signal failed to remain in the pole in *rgf1*Δ cells (Movies S1 and S2). This can be better observed in the kymographs shown in Fig 1E, where the fluorescence of Tea4 vanished from the cell cortex of the *rgf1*Δ cells in a few seconds, whereas it remained stable in control cells.

Polarity factors such as Tea1 and Tea4 localize to the cellular cortex in discrete clusters (18), which are not easily observable when taking conventional lateral cell images. To better perceive the formation of Tea4 nodes at the cell poles we used Super-Resolution Radial Fluctuations (SRRF) microscopy for “head-on” imaging of cell tips. We performed short time-lapse experiments (three minutes) at the cellular tip cortex on “head-on” wild-type and *rgf1*Δ cells (tagged with Tea4-GFP and mCherry-Atb2). In wild-type cells, Tea4 nodes deposited by MTs remained stable for an interval of time even after MT catastrophe (Fig 1F, white arrows). Moreover, clusters of Tea4 not associated with microtubules could be observed stable throughout the time-lapse, and were of the same size as those associated with MTs. However, in *rgf1*Δ cells, Tea4 dots of a similar size to those observed in the wild-type cells appeared exclusively while they were associated with MTs (Fig 1F, blue arrows). Once the MT retracted, the Tea4 cluster became gradually smaller until it eventually disappeared. These observations confirmed the results obtained with conventional microscopy methods, suggesting that Rgf1 was not necessary for the delivery of Tea4 to the cell poles, but it is required for its stable maintenance once it was released there. In addition, we ruled out a defect in the stability of the Tea4 protein in the *rgf1*Δ mutant because the half-life of the protein in the wild-type and *rgf1*Δ cultures treated with cycloheximide was comparable (Fig S1C).

During the course of these experiments, we noticed that some MTs curled around the tips in the *rgf1*Δ cells. We confirmed that the MT dynamics was not affected because the polymerization and depolymerization rates were similar in the wild-type and *rgf1*Δ cells, with a slight increase in the polymerization rate in the mutant (Fig S1D). However, the mean time that the MT stayed at the tip was ∼43 seconds in the *rgf1*^+^ cells but ∼70 seconds in the *rgf1*Δ cells (Fig 1G). Therefore, once an MT reached the cortex in the *rgf1*Δ cells, it remained there longer than in the wild-type cells. Curved MTs have already been described in *tea1*Δ cells growing at a high temperature (10). We incubated the *rgf1*Δ and *tea1*Δ mutants at 37°C for 4 hours and observed MT organization under these conditions (Fig 1H). In the *rgf1*Δ mutant, ∼10% of the cells possessed at least one MT curled around the end, similarly to the ∼12% found in the *tea1*Δ mutant, while this type of curly MT was rarely observed in the wild-type (1.7%). It is possible that a limited amount of Tea4–Tea1 at the growing pole underlies the curly phenotype seen in the absence of Rgf1.

### Rgf1 cooperates with Mod5 in Tea4 anchoring to the cellular poles

It has been shown that in cells lacking Mod5, Tea1 and Tea4 fail to accumulate to wild-type levels at the cell tips (9,16,40). Given that the *rgf1*Δ cells showed a similar defect, we analyzed the localization of Tea4 in the double mutant *rgf1*Δ*mod5*Δ compared with the single mutants. In the *mod5*Δ and *rgf1*Δ cells, most of the Tea4-GFP dots were associated with MTs, although was still some signal mainly at one of the poles (Fig 2A, yellow arrows). However, in the *rgf1*Δ*mod5*Δ double mutant, the Tea4 signal was depleted at both poles (Fig 2A, orange arrows), suggesting that Mod5 and Rgf1 share a function in Tea4 anchoring.

**Fig 2:**
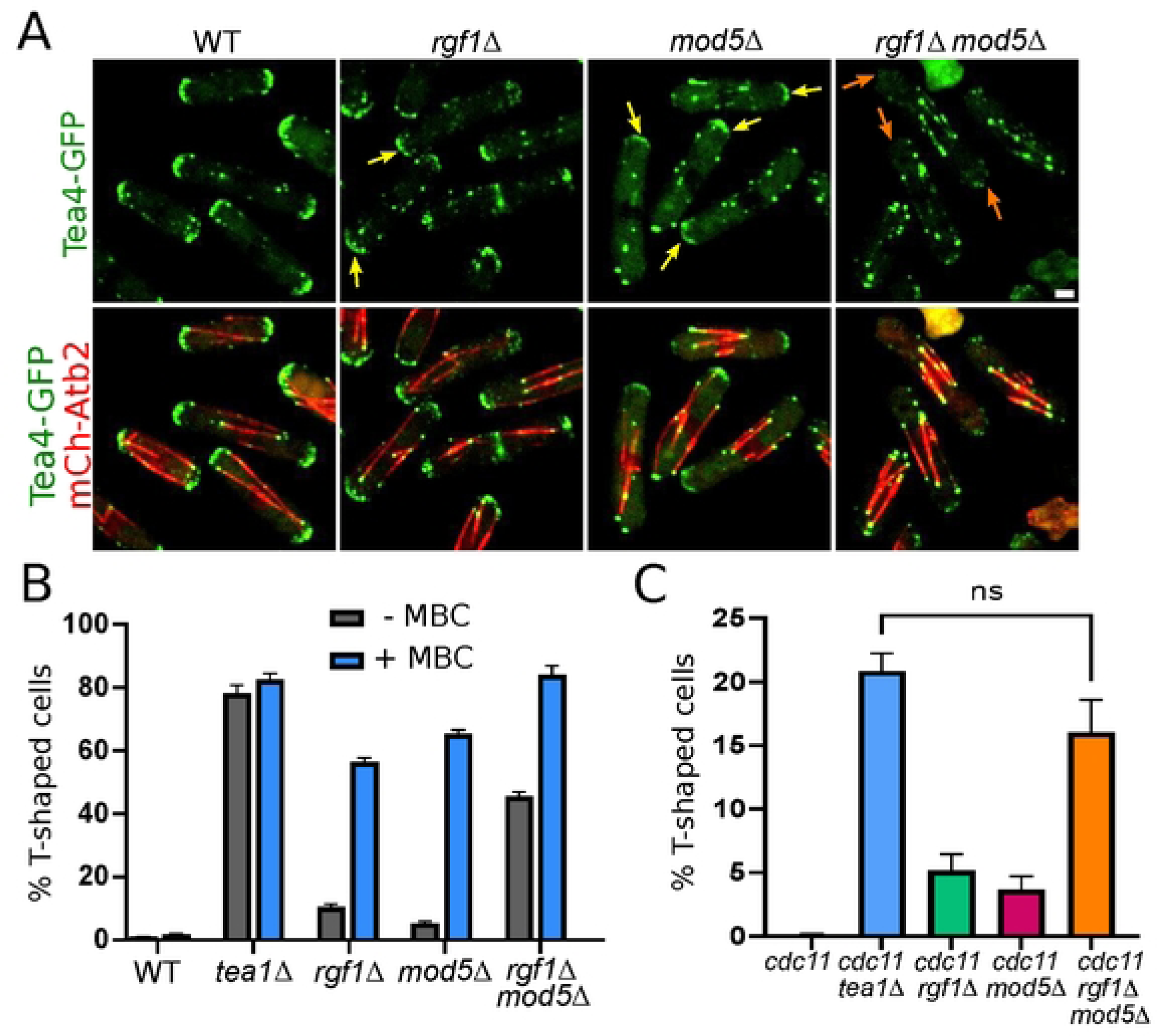
Rgf1 cooperates with Mod5 in Tea4 anchoring to the cellular poles. (A) Representative images of the indicated strains producing Tea4*-*GFP (green) and mCherry-Atb2 (red). The maximum-intensity projection of six Z-slides (0.5 µm step-size) is shown. Scale bar 2 µm. (B) The percentage of cells forming branches in the indicated strains. The cells were grown to the stationary phase for 3 days, and then were treated with DMSO (-MBC) or MBC (+MBC; 50 µg/ml). The mean ± SD of > 200 cells from three independent experiments is shown. (C) The percentage of T-shaped cells in the indicated strains after incubation for 4 hours at 36°C in YES liquid medium. Statistical significance was calculated using a two-tailed unpaired Student’s t test. ns = non-significant.

Because the localization of Tea4 is entirely dependent on Tea1 and the localization of Tea1 is partially dependent on Tea4 (9), we wondered whether the phenotype of the *rgf1*Δ*mod5*Δ cells (which mislocalize Tea4) resemble the *tea1*Δ phenotype. To this end, we scored the percentage of T-shaped cells in polarity re-establishment assays. In these assays, we first grew cells to the stationary phase for 3 days, and then diluted them in fresh medium for 3 hours. This treatment increases the penetrance of polarity mutant phenotypes, inducing T-shapes and bulges and was performed either in the presence or in the absence of the MT inhibitor methyl*-2-*benzimidazole carbamate (MBC), to prevent or allow the continuous delivery of Tea1–Tea4 to the poles by microtubules(12,16). Consistent with the “poor” localization of Tea4 in the cell cortex, the *rgf1*Δ and *mod5*Δ cells showed polarity defects when treated with MBC and marginal defects in the absence of MBC, whereas ∼80% of the *tea1*Δ cells displayed the characteristic T-shaped pattern, with or without MBC treatment (Fig 2B and S2A)(16). Interestingly, the *rgf1*Δ*mod5*Δ cells behaved similar to the *tea1*Δ mutant: reaching ∼80% of T-shaped cells with MBC and showing ∼45% even in the presence of MT (-MBC) (Fig 2B and S2A). In addition, we combined the *rgf1*Δ*mod5*Δ deletion with the temperature sensitive (ts) septation mutant *cdc11-119*. At the restrictive temperature, *cdc11-119* cells show a defect in cytokinesis, but the nuclear and growth cycles continue and cells grow at both ends after each mitosis. Presumably, after each mitosis, *cdc11-119* mutant cells must decide where to reinitiate growth. In the *cdc11-119 tea1*Δ double mutant, these events are often aberrant, leading to the formation of highly branched or T-shaped multinuclei cells (10,41). Only ∼5% of the *cdc11rgf1*Δ and *cdc11mod5*Δ cells were T-shaped after incubation for 4 hours at 36°C (Fig 2C and S2B). However, the *cdc11rgf1*Δ*mod5*Δ triple mutant showed a similar percentage of T-shaped cells (∼17%) as the *cdc11tea1*Δ double mutant (∼21%). These results indicate that Rgf1 and Mod5 collaborate to position the polarity markers at the cell tips to prevent mislocalization of growth machinery in successive cell cycles.

### Rgf1 interacts with the cell-end marker Tea4 and binds to phosphatidylinositol-4-phosphate through its PH domain

Tea1 and Tea4 reside in large protein complexes (10,19). We used different approaches to determine whether Rgf1 acts locally to retain Tea4 at the cell tips. First, we examined the *in vivo* localization of endogenous Tea4-GFP together with Rgf1-tdTomato. In newly divided and interphase cells, a subset of Rgf1 dots colocalized with Tea4 dots at cell tips, indicating a close proximity with each other (Fig 3A). Subsequently, we tested for the coprecipitation of these two proteins from yeast extracts by using epitope-tagged strains. Indeed, endogenously expressed GFP-tagged Tea4 led to the co-purification of HA-tagged Rgf1(Fig 3B). Tea1 forms a stable complex with Tea4 (9); however, we could not detect the interaction between Tea1 and Rgf1 (Fig S3). To validate these associations, we purified GST-Rgf1 from *Escherichia coli* and conjugated it with Glutathione Sepharose (GS) beads; then, we used those beads in a pull-down assay to trap Tea4-GFP or Tea1-GFP from *Shizosaccharomyces pombe* protein extracts. We detected binding of Tea4 when using Rgf1-GS beads but not with GS beads alone (Fig 3C). In addition, we could observe a slight precipitation of Tea1, which might be the result of Tea4 interaction with Tea1. We confirmed the biochemical interaction in a two-hybrid assay. Consistently, Rgf1 could interact with Tea4 but not with Tea1 (Fig 3D). Taken together, these results indicate that Rgf1 associates with the Tea1–Tea4 complex through its binding with Tea4.

**Fig 3:**
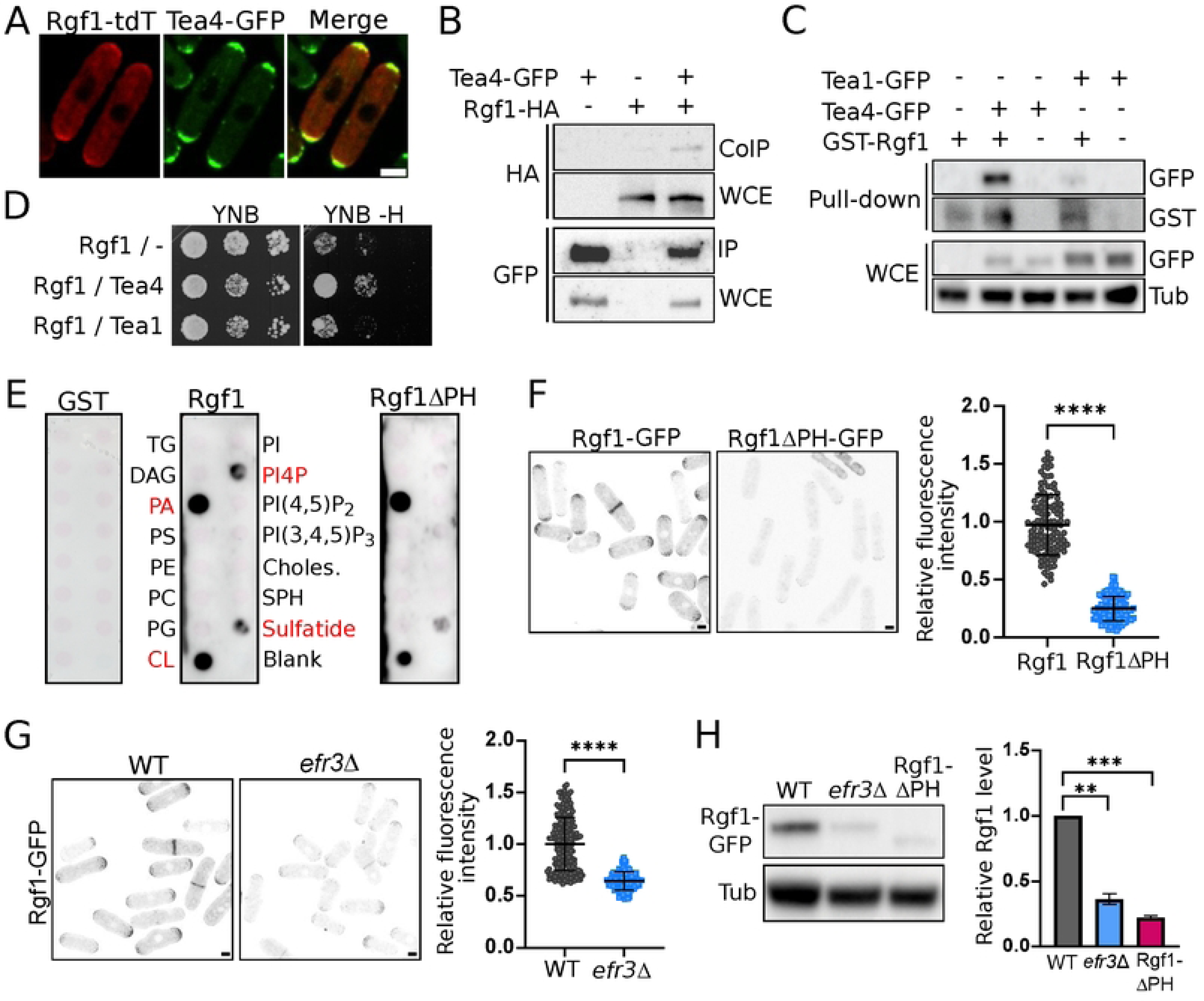
Rgf1 interacts with the cell end marker Tea4 and binds to phosphatidylinositol-4-phosphate through its PH domain. (A) Colocalization of Rgf1 and Tea4. Representative images of wild-type cells producing Tea4-GFP endogenously (green) and Rgf1-tdTomato from a plasmid under the control of its own promoter (red). The maximum-intensity projection of six Z-slides (0.5 µm step-size) is shown. (B) Coprecipitation of Rgf1 and Tea4. Cell extracts from cells producing Tea4-GFP, Rgf1-HA, or Tea4-GFP and Rgf1-HA were precipitated with GFP-trap beads and blotted with anti-HA or anti-GFP antibodies (co-immunoprecipitation and immunoprecipitation). Western blot was performed on total extracts to visualize total Tea4-GFP and Rgf1-HA levels (whole cell extracts). (C) Cells expressing Tea1-GFP or Tea4-GFP were pulled down from cell extracts with GST-Rgf1 purified from *E. coli* bound to GS-beads or with GS-beads alone, and blotted with anti-GST or anti-GFP antibodies (Pull-down). Total Tea1-GFP or Tea4-GFP levels (WCE) were visualized by western blot; tubulin was used as a loading control. (D) Two-hybrid analysis of the interaction between Tea1 (pGR135) and Tea4 (pGR106) with Rgf1 (pRZ97). The interaction was assessed on YNB plates without histidine (YNB -H). (E) Protein-lipid overlay assay. Membrane lipid strips were overlaid with 1 ug/ml of the purified GST, GST-Rgf1, and GST-Rgf1ΔPH respectively, and the interaction was detected with an anti-GST antibody. Lipids to which GST-Rgf1 showed a significant association are shown in red. Note that the interaction with PI4P disappears with GST-Rgf1ΔPH. (F) Representative images of cells producing Rgf1*-*GFP or Rgf1ΔPH-GFP. The maximum-intensity projection of six Z-slides (0.5 µm step-size) is shown. The graphic represents the mean ± SD of the relative fluorescence intensity measured at the cellular tips of Rgf1-GFP and Rgf1ΔPH-GFP (n>120). (G) Representative images of cells producing Rgf1*-*GFP in WT or *efr3*Δ mutant. The maximum-intensity projection of six Z-slides (0.5 µm step-size) is shown. The graphic represents the mean ± SD of the relative fluorescence intensity measured at the cellular tips of Rgf1-GFP (n > 120). (H) Protein extracts from cell producing Rgf1*-*GFP in the WT or *efr3Δ* mutant and Rgf1ΔPH*-*GFP were analyzed by western blot with an anti-GFP antibody to visualize Rgf1 levels. An anti-tubulin antibody was used as a loading control. The graphic represents the mean ± SD of the relative Rgf1 proteins levels from two independent experiments. Statistical significance was calculated using a two-tailed unpaired Student’s t test. ****P< 0.0001; ***P< 0.001; **P < 0.01. Scale bar 2 µm.

Next, we evaluated whether Rgf1 acts as a linker between Tea4 and the PM in the anchoring process. Rgf1 is a large (∼150 KDa) multi-domain protein, including a pleckstrin homology (PH) domain (34). PH domains could act as a “membrane-targeting device” by anchoring GEFs to phosphoinositides and directing them towards their partner GTPases on the cellular cortex (42,43). To test the ability of Rgf1 to bind different membrane lipids, we fused the Rgf1 protein to GST (without its carboxi-terminal CNH domain to purify it more easily) and purified it from bacteria. We used membrane arrays spotted with different kind of lipids to detect binding of GST-Rgf1 to lipids. As shown in Fig 3E, recombinant-purified GST-Rgf1 preferentially bound to phosphatidic acid (PA) and cardiolipin and also bound to phosphatidylinositol 4-phosphate [PI(4)P], and 3-sulfogalactosylceramide more subtly. GST-Rgf1ΔPH (additionally lacking the PH domain) could not interact with PI(4)P, but it behaved like the wild-type protein in terms of its binding to the other lipids. This result contrasts with that described for the budding yeast Rgf1 homolog, Rom2, whose PH domain specifically binds to phosphatidylinositol 4,5-bisphosphate [PI(4,5)P_2_] (44). Thus, in *S. pombe* Rgf1 might bind to the PM through the interaction between its PH domain with the phospholipid PI(4)P. Consistently, the PH domain was required for proper anchoring of Rgf1 to the cellular cortex. A GFP-tagged Rgf1ΔPH mutant showed large defects in its localization at both the poles and the septum (Fig 3F). The fluorescence detected at the *rgf1*ΔPH-GFP cell ends was ∼25% of that exhibited by the full-length protein Rgf1-GFP (Fig 3F, right).

Kathleen Gould’s group previously reported that Rgf1 displays septum localization defects in cells lacking the gene *efr3*^+^, a PM scaffold for the PI(4)P kinase Stt4. Cells lacking *efr3*^+^ do not properly position Stt4 and display altered levels of PM phosphoinositides (45). We found that the localization of Rgf1 at the cell poles was compromised in the *efr3*Δ cells (Fig 3G). Moreover, the PM binding of Rgf1 might be crucial for protein stability. We detected a 3–4-fold decrease in the protein level of Rgf1, both without the PH domain or in *efr3*Δ cells, when Rgf1 was not properly bound to the PM (Fig 3H). These results support a role for PM phospholipids in the anchoring of Rgf1 to the cellular cortex and in the maintenance of Rgf1 protein level.

### Tea4 accumulation at the cell ends depends on Rgf1 anchoring to the PM and Rho1 activation

Next, we determined whether Tea4 is bound to the cell cortex in the *rgf1*ΔPH-GFP mutant, which shows compromised Rgf1 localization. Cortical localization of Tea4 was greatly reduced (∼35%) in the mutant lacking the PH domain compared with the wild-type (P < 0.0001) (Fig 4A).

**Fig 4:**
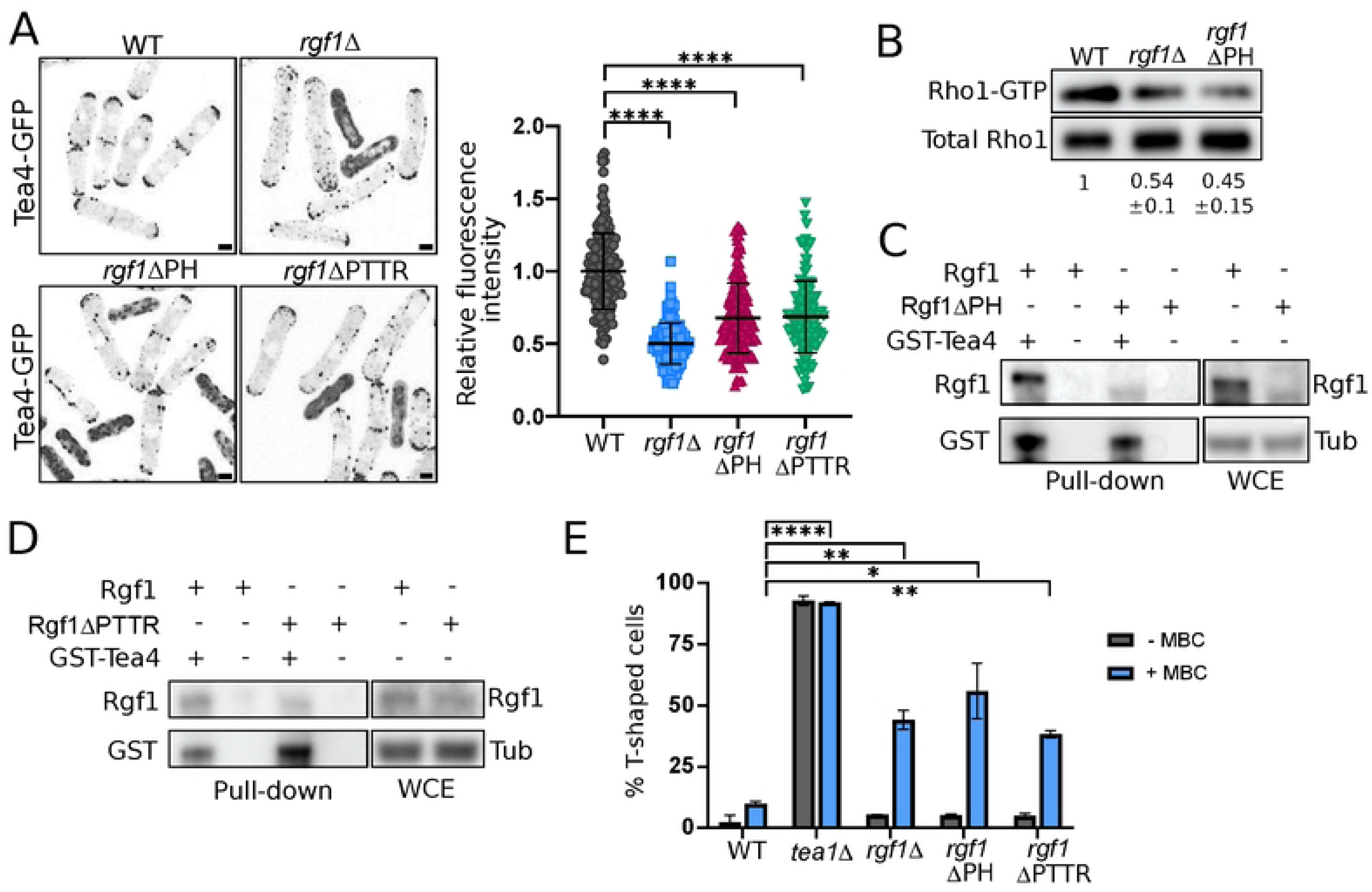
Tea4 accumulation at the cell ends depends on Rgf1 anchoring to the PM and Rho1 activation. (A) Representative images of Tea4-GFP localization in the WT, *rgf1*Δ, *rgf1*ΔPH, and *rgf1*ΔPTTR cells. The maximum-intensity projection of six Z-slides (0.5 µm step-size) of Tea4-GFP fluorescence is shown. Scale bar 2 µm. The graphic represents the mean ± SD of the relative fluorescence intensity of Tea4-GFP (n > 130) measured at the cellular tips. (B) Extracts from cells producing Rho1-HA (pREP4X-Rho1-HA) in the WT, *rgf1*Δ, and *rgf1*ΔPH cells were pulled down with GST-C21RBD and blotted against anti-HA antibody (Rho1-GTP). Total Rho1-HA was visualized by western blot (WCE). The relative units indicate the fold-differences in Rho1 levels in the mutants compared with the WT strain, with an assigned value of 1 (bottom) from three independent experiments. (C) Extracts from cells producing Rgf1-GFP or Rgf1ΔPH-GFP were pulled down with GST-Tea4 purified from *E. coli* bound to GS-beads or with GS-beads alone and blotted against anti-GST or anti-GFP antibodies (Pull-down). The total Rgf1 levels (WCE) were visualized by western blot; tubulin was used as a loading control. (D) Extracts from cells producing Rgf1-HA or Rgf1ΔPTTR-HA were pulled down with GST-Tea4 purified from *E. coli* bound to GS-beads or with GS-beads alone and blotted against anti-GST or anti-GFP antibodies (Pull-down). Total Rgf1 levels (WCE) were visualized by western blot; tubulin was used as a loading control. (E) The percentage of cells WT, *tea1*Δ*, rgf1*Δ, *rgf1*ΔPH and *rgf1*ΔPTTR cells forming branches 3 hours after release to growth after 3 days in stationary phase, in the absence and in the presence of MBC (50 µg/ml). The mean ± SD of > 200 cells from two independent experiments is shown. Statistical significance was calculated using a two-tailed unpaired Student’s t test. ****P< 0.0001; **P < 0.01; *P < 0.05.

However, in the *rgf1*ΔPH-GFP mutant the ability to join and anchor Tea4 to the cortex could be reduced due to a significant drop in the protein level compared with the Rgf1-GFP protein (Fig 3H). GTPase activation by its GEFs usually takes place when both the GTPase and the GEF are close to the PM; thus, a low level of Rgf1ΔPH-GFP at the PM could promote inefficient activation of the Rho1 GTPase. To examine whether this is the case, we analyzed the *in vivo* amount of GTP-Rho1 (active-Rho1) in the *rgf1*ΔPH cells in a pull-down assay with GST-C21RBD, the rhotekin-binding domain (previously purified from bacteria) (28). We found that the level of active-Rho1 detected in the *rgf1*ΔPH cells was similar to that seen in the *rgf1*Δ cells, and much less than the amount detected in control cells (Fig 4B). Both, the localization of Rgf1 to the PM and the activation of Rho1 were impaired in the *rgf1*ΔPH mutant; hence, we could not determine which one is behind Tea4 mislocalization. To distinguish between these two possibilities, we utilized the *rgf1-*Δ*PTTR* mutant expressing a protein without four amino acids in the RhoGEF domain. This mutant displays significantly reduced GEF activity toward Rho1,(36) while the protein remained attached to the growing end (Fig S4). Tea4-GFP was also mislocalized in the *rgf1-*Δ*PTTR* mutant (Fig 4A), indicating that the stable association of Tea4 to the membrane is dependent on Rgf1 GEF activity.

We wondered whether the Rgf1*-*Δ*PH* and Rgf1*-*Δ*PTTR* mutant proteins retained the ability to bind Tea4 *in vitro*. Both proteins proficiently bound Tea4 in an *in vitro* pull-down assay (Fig 4C and D). *In vivo*, we analyzed the percentage of T-shaped cells after re-entry from the stationary phase to fresh medium. With MBC treatment, the percentage of T-shaped cells was approximately 50% for the *rgf1*Δ mutant; this percentage was similar for the *rgf1-*Δ*PH* (∼55%) and *rgf1-*Δ*PTTR* (∼35%) mutants and much higher than for the wild-type strain (∼5%) (Fig 4E). Taken together, these results indicate that the localization of Rgf1 to the PM and its ability to activate Rho1 are closely linked: Both are required to maintain Tea4 at the cell tips and to preserve the growth pattern after refeeding. Given that the Rgf1-Δ*PTTR* protein localized at the growing end, where it is incompetent for Rho1 activation, GEF activity appears to be more critical than GEF localization regarding Tea4 maintenance at the poles. Thus, Rho1 could act by promoting the formation of a stable critical mass of Tea4 in the cell cortex that is essential to activate pole-restricted growth.

### Rgf1 is part of actin-dependent machinery that signals growth poles in the absence of the Tea1–Tea4 complex

As we have described above, *tea1*Δ cells experience an exacerbated loss of polarity when subjected to nutritional stress while keeping their cylindrical shape in normal conditions. We wondered why this occurs and how cells recognize their poles in the absence of the classical Tea markers. Since Tea4 localization is entirely dependent on Tea1(9) we utilized *tea1*Δ mutant to eliminate both markers at the poles. It has been proposed that the high number of T-shaped cells observed in the *tea1*Δ mutant in refeeding experiments (Fig 2B and 4E) is due to a transient depolarization of the actin cytoskeleton (46). To address this issue, we confirmed the disorganization of the actin cytoskeleton in wild-type cells grown stress treatments exposure to KCl, sorbitol or heat, which promoted actin depolarization also induced branching in the *tea1*Δ mutant for 3 days in liquid medium (Fig 5A). In addition, (Fig S5A and B) (11,47). We thought that if the mechanism that keeps the identity of the growth sites in the absence of Tea1 is lost because of transient actin depolarization, then treating *tea1*Δ cells with latrunculin A (LatA), which prevents the polymerization of filamentous actin, should increase the number of branched cells. As expected, 75% of the *tea1*Δ cells showed polarity defects during recovery from the LatA treatment, compared with ∼2% of the wild-type and untreated *tea1*Δ cells (Fig 5B). Thus, a properly polarized actin cytoskeleton is required to position growth sites at opposite cell poles in the absence of Tea1**–**Tea4 markers.

**Fig 5:**
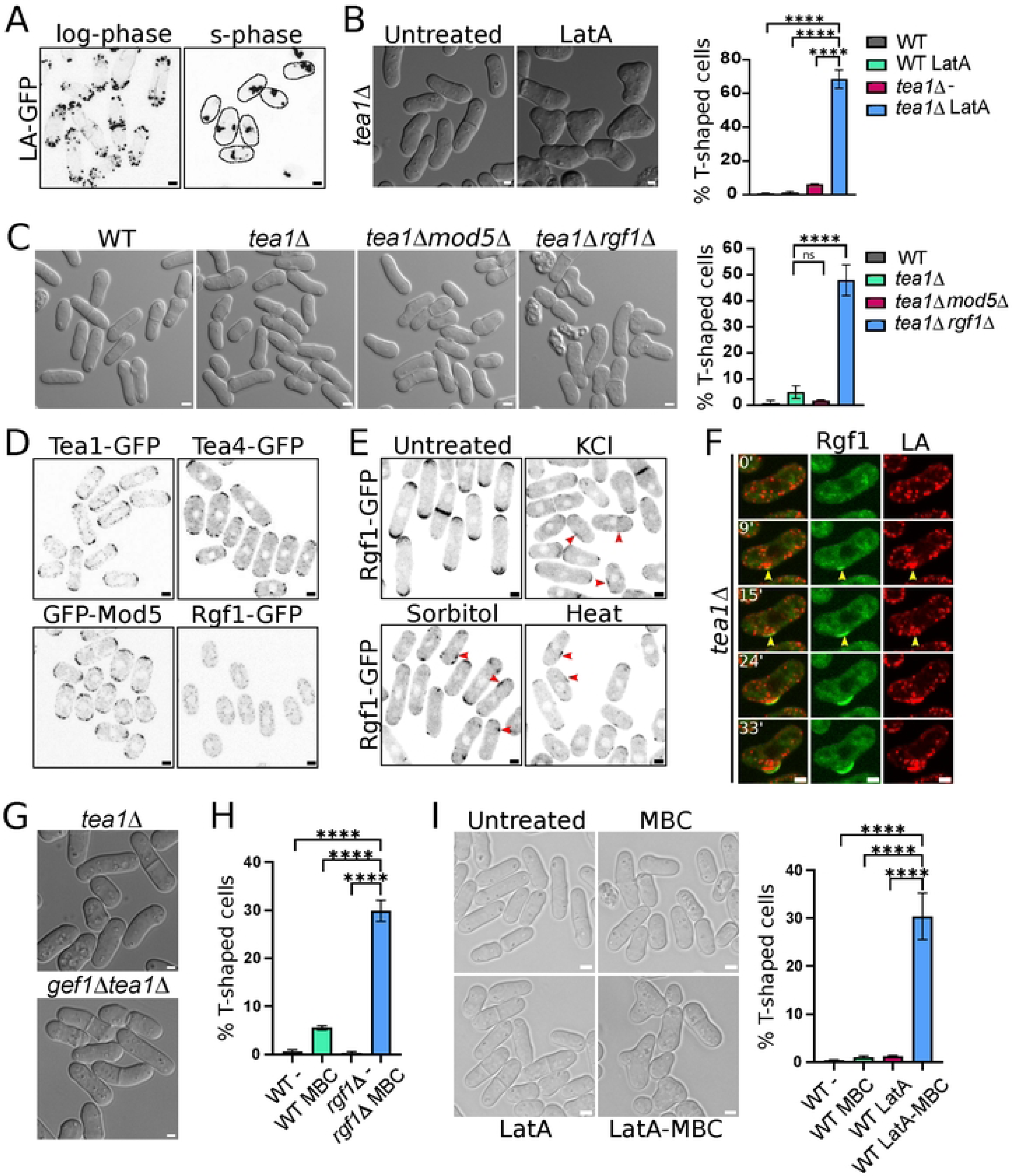
Rgf1 is part of an actin-dependent machinery that signals growth poles in the absence of the Tea1–Tea4 complex. (A) Images of LifeAct-GFP (actin) localization in the WT cells growing in liquid medium in the log-phase or after 3 days in the stationary phase (s-phase). The maximum-intensity projection of six Z-slides (0.5 µm step-size) of fluorescence is shown. (B) Morphology and quantitation of the T-shaped wild-type and *tea1*Δ cells treated for 2 hours with DMSO (untreated) or 50 µM of Latrunculin A (LatA) and then washed to allow growth for 3 hours. The graph represents the mean ± SD of > 200 cells from three independent experiments. (C) Morphology and quantitation of the T-shaped cells in the WT, *tea1*Δ, *tea1*Δ *mod5*Δ, and *tea1*Δ *rgf1*Δ cells grown to log phase in YES liquid medium at 28°C. The graph represents the the mean ± SD of > 500 cells from two independent experiments. (D) Representative images of Tea1-GFP, Tea4-GFP, GFP-Mod5 and Rgf1-GFP localization in wild-type cells in the stationary phase after 3 days of growth in liquid medium. The maximum-intensity projection of six Z-slides (0.5 µm step-size) of fluorescence is shown. (E) Rgf1-GFP localization in WT cells growing in liquid medium untreated or treated with KCl 0.6M, sorbitol 1.2M or 37°C (heat) for 1 hour. The maximum-intensity projection of six Z-slides (0.5 µm step-size) of fluorescence is shown. The arrowheads point lateral accumulation of Rgf1-GFP. (F) LifeAct-mCherry (actin in red) and Rgf1-GFP localization (green) in *tea1*Δ cells treated with KCl 0.6M for 1 hour and then washed and allowed to grow without stress for the indicated times. The maximum-intensity projection of four Z-slides (0.6 µm step-size) of fluorescence is shown. The arrowheads point to lateral accumulation of Rgf1-GFP and actin (LA). (G) Morphology of the *tea1*Δ and *gef1*Δ*tea1*Δ cells growing in YES liquid medium. (H) Quantitation of the T-shaped cells in WT and *rgf1*Δ cells treated with DMSO (-) or with MBC (50 µg/mL) for 4 hours. The graph represents the mean ± SD of > 200 cells from three independent experiments. (I) Morphology and quantitation of the T-shaped cells in wild-type cells treated with DMSO (WT -), MBC 50 µg/mL (WT MBC) for 4 hours, 50 µM of LatA for 2 hours and then washed and allowed to grow with DMSO for 4 hours (WT LatA) or first treated with LatA, washed, and then treated with 50 µg/mL of MBC for 4 hours (WT LatA-MBC). The graph represents the mean ± SD of > 200 cells from three independent experiments. Statistical significance was calculated using a two-tailed unpaired Student’s t test. ****P < 0.0001; ns = non-significant. Scale bar 2 µm.

Given that Rgf1 is required for actin re-organization during NETO (28), we reasoned that the absence of Rgf1 in the *tea1*Δ background could uncover polarity defects that would otherwise remain undetected. This was indeed the case; 50% of the *tea1*Δ*rgf1*Δ cells in unperturbed conditions were T-shaped compared with fewer than 5% of the *tea1*Δ and *tea1*Δ*mod5*Δ cells (Fig 5C). Thus, in the absence of Rgf1 and polarity markers (Tea1–Tea4), stresses that induce actin depolarization were not necessary for branching to occur. Moreover, Rho1 activation was required to maintain polarity, evidenced by the high number of T-shaped cells observed in the *tea1*Δ*rgf1*ΔPTTR mutant (Fig S5C). Therefore, in the absence of Tea1, Rgf1– Rho1 probably mark the poles through an actin-dependent mechanism.

To understand why the *tea1*Δ*rgf1*Δ null cells behaved alike *tea1*Δ stressed cell, we studied the localization of Rgf1 under situations that depolarize the actin cytoskeleton. Rgf1-GFP localization to the cell tips was lost quickly in cells treated with LatA, but was unaffected in cells treated with MBC (Fig S5D). Thus, we reasoned that because actin is depolarized in quiescent cells, Rgf1-GFP should behave similarly. Accordingly, Rgf1 was missing from the cell periphery in stationary phase cells, while Tea1, Tea4, and Mod5 (which depend on MTs to reach the poles), remained polarized in the same experiment (Fig 5D) (10,11,16). Moreover, Rgf1 disappeared from the cell tips under osmotic or heat stress but was observed in lateral patches (Fig 5E). Then, we followed actin reorganization and Rgf1 localization during recovery from osmotic stress in wild-type and *tea1*Δ cells. After relieving the stress, wild-type cells quickly re-concentrated both actin and Rgf1 at the poles (Fig S5E and Movie S3). However, in *tea1*Δ cells actin and Rgf1 frequently localized at the lateral cortex precisely at points where a branch began to grow (Fig 5F and Movie S4).

To find out whether other proteins of the growth machinery also define the growth sites or if it is a specific characteristic of Rgf1, we used a mutant lacking *gef1*. Gef1 is a GEF of Cdc42 involved in polarized growth and undergoes a similar relocation from poles to lateral patches under stress, where it is required to activate Cdc42 (48,49). Interestingly, the *gef1*Δ*tea1*Δ double mutant displayed a similar percentage of T-shaped cells as the *tea1*Δ mutant, indicating that in the absence of Tea1, Gef1 is not necessary for marking the poles, in contrast to Rgf1–Rho1 (Fig 5G and S5F).

Taken together, these results indicated that Rgf1 localization at the cell tips depends on a properly polarize actin cytoskeleton but it is independent on MTs. Thus, both actin concentration and Rho1 activation at the poles are necessary events to define these locations as growth sites.

### Tea1–Tea4-MTs and Rgf1-Rho-actin define two parallel pathways to restrict growth to the cell tips

Because stresses that induce actin disorganization and Rgf1 delocalization at the poles increase the percentage of T-shaped cells in the *tea1*Δ mutant (Fig 5D and S5A), we wondered whether chemical ablation of the pathway that delivers Tea1 and Tea4 to the cell tips would yield a similar result. When we treated the *rgf1*Δ mutant with MBC, ∼30% of the cells exhibited a branched phenotype compared with ∼5% of the wild-type cells (Fig 5H and S5G). Therefore, in the absence of Rgf1 (which lacks the “actin-dependent signal” necessary to recognize the polarity growth zones) and MTs, the imposition of stress is not necessary for branching to occur. The previous results suggest that there could be two parallel pathways for positioning the growth poles: one dependent on MTs and the Tea1–Tea4 markers, and another dependent on actin and Rgf1– Rho1. To mimic the elimination of both signaling pathways by chemical treatment, we treated wild-type cells in the log phase with LatA for 2 hours (to block actin-dependent signaling), washed them, and exposed them to MBC for 4 hours (to remove MTs) to analyze the number of T-shaped cells. Only after the treatment with both, LatA and MBC, ∼35% of the cells showed branches (Fig 5I).

Our results indicate that Rgf1–Rho1 and Tea1–Tea4 are part of the same complex, with similar functions to delimit the sites of polarized growth of *S. pombe*. We propose that Rgf1 and Rho1 could activate an actin-dependent pathway that instructs cells to grow at the tips regardless of the classical polarity markers.

## DISCUSSION

Cross-talk between microtubules (MT) and the actin cytoskeleton is crucial for various cellular processes, including asymmetric cell division, the establishment of cell growth zones, and cell migration. In fission yeast, MTs deliver the Tea1–Tea4 complex to the cell tips, where actin concentrates to promote growth. Tea1–Tea4 act as end markers, defining the organization of cell-growth zones and, consequently, the direction of growth. While it is known that these polarity markers are not essential for organizing a growth zone, they become critical for placing the growth zone correctly, especially under stress conditions. However, certain questions remain unanswered, such as how and why the Tea1–Tea4 complex remains linked to the plasma membrane (PM) once the growth direction is established and how cells mark their tips in the absence of Tea1. In the present study, we addressed some of these questions by demonstrating that Rgf1 functions as a molecular link between the Tea1-Tea4 complex and the PM. Through Rho1 activation, Rgf1 stabilizes Tea4 at the cell ends, promoting its accumulation. Additionally, we described an alternative actin-dependent mechanism, driven by Rgf1 and Rho1, for marking the poles independently to the known MT-and Tea-dependent pathway.

### Rgf1 (Rho1 GEF) activity toward Rho1 is required for stable accumulation of Tea4 at the cell ends

Failure to accumulate Tea4 at the cell cortex in the *rgf1*Δ cells is independent of the movement of Tea4 riding on the tips of polymerizing MTs. Wild-type and *rgf1*Δ cells exhibited similar rates of movement for Tea4-GFP dots from the middle of the cell to the cell ends (Fig S1D). However, once the Tea4-GFP dots had reached the cell ends, their fluorescence faded in the *rgf1*Δ cells. The refill mechanism responsible for keeping Tea4 stacked at the growing pole depends on correct binding of Rgf1 to the PM and on Rgf1 catalytic activity toward Rho1. We provided evidence that Rgf1 physically binds the polarity marker complex through its interaction with Tea4 (Fig 3B-D). Moreover, Rgf1 binds to PM phospholipids, likely through its PH domain interacting with membrane PI4P. The physical association between Rgf1 and Tea4 is crucial for the localization of Tea4 and Tea1 at the cell poles. Indeed, a mutant of Rgf1 lacking the membrane-binding domain, which binds Tea4 *in vitro*, exhibited similar polarity defects as the null mutant (Fig 4E). In the *rgf1*ΔPH mutant, the reduced activation of Rho1 (Fig 4B) is coupled to protein instability (Fig 3H) (probably because it is not properly bound to the PM) which makes the interpretation of this result challenging. Additionally, a catalytic mutant (*rgf1-*ΔPTTR) also shows impaired Tea4 anchoring. The Rgf1*-*ΔPTTR protein binds Tea4 *in vitro*, localizes to the growing end and is catalytically deficient (36), indicating that Rho1 activity is required to maintain a the Tea1-4 complex stable at the pole. Furthermore, the tandem Rgf1–Rho1 also affects Tea1 functions, promoting MT catastrophe once they reach the cell pole or restricting growth sites after refeeding (Figs 1G-H and 2B). To maintain polarity markers stably at the poles, Rgf1 functions together with Mod5. The loss of Tea4 localization at the poles is partial in the *rgf1*Δ and *mod5*Δ single mutants and complete when both mutations are combined, causing a defect in polarity even in cells with a continuous supply of Tea1-Tea4 (Fig 2B and C). It is likely that Mod5 and Rgf1 anchor the Tea1-4 complex to the PM through different proteins: Mod5 interacts with Tea1 (17), while Rgf1 interacts with Tea4. This mechanism ensures double anchoring of the Tea1–Tea4 complex to the PM; thus, when one of these connections is lost, the other remains available. Only when both are missing do the polarity markers completely lose their connection with the PM, mimicking the behavior of a the *tea1Δ* mutant.

### Rgf1–Rho1 are involved in defining the growth sites

Another question that must be addressed is how cells detect their growth sites in the absence of Tea markers. Here we proposed that actin and Rho1 activation by Rgf1 are behind this process. We observed two ways to induce the formation of branched cells in the *tea1*Δ background: one in the presence of certain stresses, “the stress pathway” and the other in the absence of *rgf1*, “Rgf1 depletion” (Fig 5C and S5B). The loss of polarity induced by “the stress pathway” also leads to disorganization of the actin cytoskeleton and, consequently, Rgf1 delocalization. Therefore, both pathways lead to insufficient activation of Rho1 at the poles.

When the *tea*1Δ cells recover from stress, Rgf1 and actin appear simultaneously at sites were ectopic growth occurs. The interdependence of actin and Rgf1 localization (Fig S5D) (28) suggests a positive feedback loop between Rho1 activation and actin organization at the growth sites. Therefore, Rgf1–Rho1 would act as Tea markers, defining growth sites without directly promoting growth, given that the *tea1*Δ*rgf1*Δ double mutant retains the ability to grow, albeit at the wrong places. Interestingly, in the absence of Tea1 and Gef1 (Cdc42 GEF), which is also involved in polarized growth and relies on actin for proper localization (48–50), cells do not form branches, suggesting that is the activation of Rho1 and not that of Cdc42 which specifically defines the growth sites.

The interaction between Tea4 and Rgf1–Rho1 indicates their involvement in the same complex, with each protein essential for the accumulation of the other at the poles. Rgf1 together with Mod5 acts to link Tea4 to the membrane, and Tea1-4 assists in returning Rgf1 to the poles after stress-induced actin disorganization (Fig 5F and S5E), establishing a functional link between MTs and actin cytoskeletons (Fig 6A).

**Fig 6:**
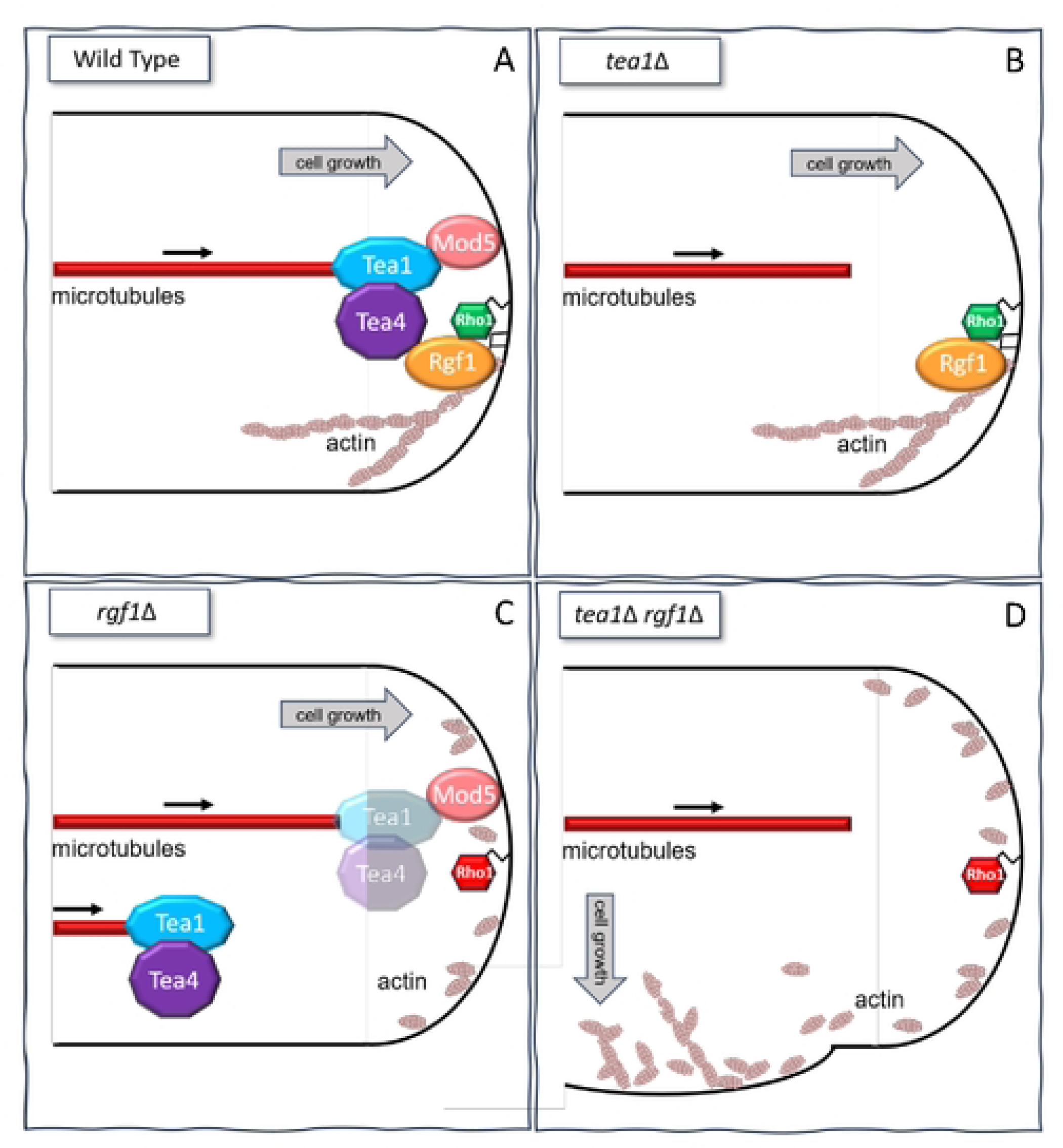
Rgf1-Rho1 functions as a molecular link between Tea4 and the PM and marks the growth sites in an actin-dependent manner. (A) In wild-type cells MT deliver Tea1 and Tea4 to the cell poles, where they bind to the membrane through their interaction with Mod5 and Rgf1, respectively. Rgf1, in turn, interacts with the PM due to its affinity for the phospholipid PI4P and activates Rho1, promoting proper actin cytoskeleton polarization at the cellular tips. (B) Cells lacking Tea1 still recognize the cell poles correctly because Rgf1 is localized in these regions, where it activates Rho1, thus allowing the maintenance of polarized actin. (C) In *rgf1*Δ cells Tea1-Tea4 partially disappears from the pole, although a remnant remains attached to Mod5. Rho1 would be inactive, leading to actin cytoskeleton disorganization. However, the continuous supply of Tea1–Tea4 from MTs persists, allowing the cells to grow correctly. (D) When cells lack *tea1* and *rgf1*, they lose both pathways that allowed them to distinguish the tips. First, there is no continuous supply of polarity markers towards the cell ends by the MTs that mark the growth sites. Second, Rgf1 is not at the pole to activate Rho1, leading to actin disorganization. The elimination of both pathways causes the cells to direct their growth towards incorrect locations, where growth factors probably accumulate.

We showed that Rgf1 and Rho1 are required to maintain an actin-dependent signal that preserves the identity of the cell poles, which becomes evident in the absence of the Tea markers. This would explain why the *rgf1*Δ*tea1*Δ double mutant forms T-shaped cells in the log phase, without stresses or refeeding treatments. We propose that there are two different pathways to choose the growth sites under different environmental conditions: the canonical pathway dependent on MT and Tea1–Tea4 and, a novel pathway dependent on actin and Rgf1– Rho1. Various combinations of mutants and/or chemical ablation of one component from each pathway (Tea1–Tea4–MT and Rgf1–Rho1–actin) at once, leads to comparable outcomes. For example, chemical actin depolymerization in the *tea1*Δ cells induces branching (Fig 5B), whereas treatment of the *rgf1*Δ cells with MBC to prevent the constant supply of Tea markers generates T-shaped cells as well (Fig 5H). Furthermore, wild-type cells treated to transiently depolarize actin and subsequently prevented for refeeding of polarity markers to the tips (via MT de-polymerization) also experience difficulties in detecting the cell poles (Fig 5I). Thus, cells would have different pathways to maintain their cylindrical morphology even if one of them is challenged by internal or external conditions. When the continuous supply of Tea1 from MT is disrupted (Fig 6B), cells could use an additional cue provided for a different cytoskeletal polymer, actin. Similarly, when actin becomes disorganized, for example, in the transition from monopolar to bipolar growth (NETO) or in *rgf1*Δ mutant (defective in NETO), cells would count on polarity markers transported by MTs (Fig 6C). Only when both, the actin and MT cytoskeletons are compromised, fission yeast cells lose their cylindrical shape (Fig 6D). This sophisticated regulation highlights the importance of maintaining cell morphology throughout the cell cycle and under changing environmental conditions.

## MATERIALS AND METHODS

### Media, Reagents and Genetics

*S. pombe* strains were streaked on plates of complete yeast growth medium (YES), or selective medium (EMM) supplemented with the appropriate requirements (51), and incubated at 28°C until colonies formed. For each biological replicate, a single colony was used to inoculate 5 mL of the respective liquid media. Cultures were incubated at 28°C overnight with shaking (200 rpm). Each overnight culture was subsequently used as a seed culture to inoculate fresh media. Fresh cultures were next grown at 28°C, 200 rpm, to OD 600 = 0.5-0.6 at the time of harvest. Crosses were performed by mixing the appropriate strains directly on sporulation medium plates. Recombinant strains were obtained by tetrad analysis or the “random spore” method. For overexpression experiments using the nmt1 promoter, cells were grown in EMM containing 15 µM thiamine up to the logarithmic phase. Then, the cells were harvested, washed three times with water, and inoculated in fresh medium (without thiamine) at an optical density at 600 nm (OD_600_) of 0.01 for 18-20 hours. Two-hybrid interaction was tested with YNB medium lacking histidine in *Saccharomyces cerevisiae* strain AH109 (Clontech).

### Plasmid and DNA manipulations

Plasmids used in this study are listed in Key resources table (Recombinant DNA). pREP4x-HArho1 (with thiamine-repressible *nmt1* promoter) and pGEX-C21RBD plasmids (rhotekin-binding domain) kindly provided by Pilar Pérez (Instituto de Biología Funcional y Genómica, Salamanca, Spain) were used to detect Rho1-GTP levels. To express proteins in *Escherichia coli* we used a pGEX-2T plasmid that contains a GST to tagged genes al 5’. pGEX-*rgf1* (pGR152) and pGEX-*tea4* (pGR129) were made by inserting the entire ORF of *rgf1* (without introns) or *tea4* in frame into pGEX-2T and purified for be used in pull-down assays. To perform lipid strips assays we constructed the plasmids pGR128 that contains the ORF of *rgf1* (from aminoacid 1-922) without the last 428 aminoacids containing the CNH domain, and pGR138 that contains the ORF of *rgf1* (from aminoacid 1-722) without the last 578 aminoacids containing the PH and CNH domains. For two-hybrid experiments we constructed the plasmids pGAD-*tea1* (pGR135), pGAD-*tea4* (pGR106), and pGBK-*rgf1* (pRZ97) where the entire ORF of the corresponding gene was inserted in frame into the pGADT7 (GAL4 activation domain) or pGBKT7 (GAL4 binding domain) plasmid (Clontech).

### Protein extracts and immunoblot analysis

*S. pombe* cultures (5 mL) at an OD_600_ of 0.5 were pelleted just after the addition of 10% TCA and washed in 20% TCA. The pellets were resuspended in 100 μl 12.5% TCA with the addition of glass beads and lysed by vortexing for 5 min. Cell lysates were pelleted, washed in iced acetone, and dried at 55°C for 15 min. Pellets were resuspended in 50 μl of a solution containing 1% SDS, 100 mM Tris–HCl (pH 8.0), and 1 mM EDTA. Samples were electrophoretically separated by SDS-PAGE (4–15% MiniProtein Gel, BioRad) and immunodetected with anti-GFP (Living Colors, RRID:AB_10013427) and anti-mouse (Bio-Rad, AB_11125547) antibodies. As a loading control, we used monoclonal antitubulin antibodies (Sigma, RRID:AB_477579).

### Immunoprecipitations and pull-down assays

Immunoprecipitation assays were performed as described previously with some modifications (52). Briefly, logarithmic cultures were pelleted and re-suspended in lysis buffer (10 mM Tris– HCl pH 7.5, 150 mM NaCl, 0.5 mM EDTA, 0.5% NP40, containing 100 μM PMSF, leupeptin, and aprotinin) and lysed in a cryogenic grinder. Lysates were centrifuged for 5 min at 6000 g, and then 10 μL of GFP-Trap magnetic beads (Chromotek) was added to the supernatants an incubated for 1 hour at 4°C. Immunoprecipitates were washed three times with dilution buffer (10 mM Tris–HCl pH 7.5, 150 mM NaCl, and 0.5 mM EDTA). Proteins were released from immunocomplexes by boiling for 5 minutes in sodium dodecyl sulfate (SDS) loading buffer. Samples were separated by SDS–polyacrylamide gel electrophoresis (4–15% Mini-Protean TGX gels, Bio-Rad) and detected by immunoblotting with polyclonal anti-HA (Roche, RRID:AB_514506) or anti-GFP antiserum (Living Colors, RRID:AB_10013427). For pull-down assays, first GST-tagged Rgf1, Tea4, or C21RBD (rhotekin-binding domain to detect Rho1-GTP levels) was purified from *E. coli* (BL21). The fusion proteins were produced by adding 0.5 mM IPTG at 18°C overnight (or 3 hours at 28°C for C21RBD). Cells were sonicated and proteins immobilized on glutathione-Sepharose (GS) 4B beads (GE-Healthcare). After incubation for 1 hour, the beads were washed several times, and the bound proteins were analyzed by SDS-PAGE and stained with Coomassie brilliant blue. Pull-down assays were performed as described previously (53). In brief, extracts from the indicated strains were obtained by using 500 μL of lysis buffer (50 mM Tris–HCl pH 7.5, 20 mM NaCl, 0.5% NP-40, 10% glycerol, 0.1 mM dithiothreitol, and 2 mM Cl_2_Mg, containing 100 μM PMSF, leupeptin, and aprotinin) and lysed in a cryogenic grinder. Cell extracts (2–3 mg of total protein) were incubated with 2–10 μg of GST-tagged protein coupled to GS beads for 2 hours, washed four times with lysis buffer, and blotted with an anti-HA or anti-GFP antibody. Protein levels in whole-cell extracts (80 μg of total protein) were monitored by western blot. Tubulin and GST were used as loading controls.

### Lipid strip overlay assays

Lipid strip overlay assays were performed as described previously (54) using lipid strip membranes (p-6002, Echelon). Strips were blocked with 3% fatty acid–free bovine serum albumin (BSA; Sigma) in TBS-T (10 mM Tris–HCl pH 8.0, 150 mM NaCl, and 0.1% Tween-20; TBST-BSA) at room temperature for 1 hour and then incubated for 2 hours with 1 μg/mL GST, GST-Rgf1, or GST-Rgf1ΔPH in TBST-BSA. The strips were then washed three times with 5 mL of TBST-BSA and incubated with anti-GST horseradish peroxidase–conjugated antibody (GE Healthcare, RRID:AB_771429) diluted in TBST-BSA. Bound protein was detected using an enhanced chemoluminescence detection kit (BioRad). GST, GST-Rgf1, and GST-Rgf1ΔPH were expressed in *E. coli* (BL21) and purified with GS beads (GE-Healthcare) according to the manufacturer’s instructions as described above. Once attached to the beads, they were washed and eluted with 200 μL of elution buffer (100 mM Tris–HCl pH 8.0, 120 mM NaCl) with 20 mM of L-glutathione reduced (Sigma) freshly added for 30 minutes at 4°C. Aliquots were frozen in liquid nitrogen with 15% of glycerol and stored at -80°C until use.

### Microscopy

Wet preparations were observed with an Andor Dragonfly 200 Spinning-disk confocal microscope equipped with a sCMOS Sona 4.2B-11 camera (Andor) and controlled with the Fusion 2.2 acquisition software; or a Personal Deltavision (Applied Precision, LLC) microscope equipped with a CoolSNAP HQ2 camera (Photometrics) and controlled with softWoRx Resolve 3D. Depending on the experiment, a single focal plane at the centre of the cell or a stack of 4–6 images covering the entire volume of the cell (Z-series) with a spacing of 0.5–0.6 μm were captured, and the maximum projection was generated.

To analyze protein dynamics, time-lapse experiments were performed with cells in μ-Slide 8-well (Ibidi) coated with soybean lectin (Sigma Aldrich) and imaged at the indicated times using an Andor Dragonfly Spinning-disk confocal microscope. We used the ImageJ 1.53t software to calculate the relative fluorescence intensity of each protein at the cell tips. We have designed an ImageJ macro to automatically select the fluorescence regions (ROI manager) of the cell poles and to measure the fluorescence intensity of 40–130 cells. We used the integrated density value of each tip of the different strains versus the average of the integrated density of the wild-type strain to calculated the relative fluorescence levels. To create kymographs, we drew a line from the cell’s centre to the pole along a microtubule on a time-lapse movie and utilized the KymographBuilder plugin of the ImageJ software. For super-resolution radial fluctuations (SRRF) images of Tea4-GFP (green) and mCherry-Atb2 (red) in “head-on” cell tips, cells were mounted onto a μ-Slide pre-coated with lectin. Bound cells found to be frontally arrayed (“head-on”) were visually selected for imaging using a the SRRF module of the Andor Dragonfly Confocal microscope. Fifty repetitions of one focal plane were taken for each time point. The time projection of the three images at different time points is shown to follow Tea4 cluster movement. Calcofluor white (Blankophor BBH, Bayer Corporation) staining was performed by adding 1 μL of a stock solution (2.5 mg/mL) to 500 μL of samples for 20 seconds, followed by a wash with phosphate–buffered saline (PBS).

### Quantification and statistical analysis

Statistical analyses and graphs were generated using GraphPad Prism Software version 9.5.1. To compare two conditions, a two-tailed unpaired Student’s t-test was applied to determine statistical significance (as detailed in the Fig legends). P < 0.05 were considered significant. The graphs show the mean ± standard deviation of the indicated data. Asterisks represent the following: *P < 0.05 **P < 0.01; ***P < 0.001 ****P < 0.0001.

## Author contributions

Conceptualization, P.G., and Y.S.; Methodology, P.G.; Investigation, P.G., R.C. and T.E.; Writing – Original Draft, P.G.; Writing – Review & Editing, P.G., and Y.S.; Funding Acquisition, Y.S.; Supervision, P.G. and Y.S.

## Acknowledgments

We thank J.C. Ribas, Phong T. Tran, Sophie Martin, Sergio Rincón, Ken Sawin, Paul Nurse, James Moseley, Kathleen L. Gould, César Roncero, Henar Valdivieso, Sergio Moreno and Pilar Pérez for sharing strains and plasmids. We also wish to thank Javier Encinar del Dedo and Sergio Rincón for their very helpful comments on the manuscript. We are grateful to Jesús Pinto (IBFG bioinformatics facility) for ImageJ macro used for fluorescence quantification.

## Supporting Information

**S1 Table. List of yeast strains used in this study.**

**S2 Table. List of plasmids used in this study.**

**S1 Fig. Rgf1 is required for proper localization of the Tea1-Tea4 complex at the cell tip.**

(A) Wild-type cells expressing rgf1-GFP were stained with Calcofluor white (20 μg/mL) to spot areas of growth. The arrows indicate the localization of Rgf1 at the growing tip in monopolar cells. (B) Quantitation of the number of Tea4 dots associated to the MTs in WT and rgf1Δ strains per cell. The mean ± SD of > 80 cells is shown. (C) The wild-type and rgf1Δ cells expressing tea4-GFP were treated with the translation inhibitor cycloheximide (CHX, 100 μg/mL) for the indicated times. Proteins were visualized by western blot with antibodies against GFP (Tea4) or tubuline (Tub), as a loading control (upper). The graphic represents the quantification of Tea4 levels at different times (hours) after the treatment relative to time 0, which was assigned a value of 1 (bottom). (D) The graphics show the MT polymerization (left) and depolymerization (right) rates in wild-type and rgf1Δ cells. The mean ± SD of > 75 cells is shown. Statistical significance was calculated using two-tailed unpaired Student’s t test. *P < 0.05; ****P < 0.0001; ns= non-significant.

**S2 Fig. Rgf1 cooperates with Mod5 in Tea4 anchoring to the cellular poles.**

(A) Morphology of the wild-type, rgf1Δ, mod5Δ and rgf1Δmod5Δ strains after refeeding treatment (-MBC). Scale bar 2 μm. (B) Cell morphology of the indicated strains after 4 hours at 36°C in YES liquid medium. Scale bar 2 μm.

**S3 Fig. Rgf1 interacts with the cell end marker Tea4 and binds to phosphatidylinositol-4-phosphate through its PH domain.**

Coprecipitation of Rgf1 and Tea1. Cell extracts from cells producing Tea1-GFP, Rgf1-HA, and Tea1-GFP and Rgf1-HA were precipitated with GFP-trap beads and blotted with anti-HA or anti-GFP antibodies (co-immunoprecipitation and immunoprecipitation). Western blot was performed on total extracts to visualize total Tea1-GFP and Rgf1-HA levels (whole cell extracts). S4 Fig. Tea4 accumulation at the cell ends depends on Rgf1 anchoring to the PM and Rho1 activation. Representative images of Rgf1-GFP and Rgf1ΔPTTR-GFP localization. Cells were stained with calcofluor white (20 μg/mL) to spot areas of growth. Scale bar 2 μm.

**S5 Fig. Rgf1 is part of actin-dependent machinery that signals growth poles in the absence of Tea1–Tea4 complex.**

(A) Representative images of LifeAct-GFP (actin) localization in cells untreated or treated with KCl 0.6 M, sorbitol 1.2 M, or 37°C (heat) for 1 hour. The maximum-intensity projection of six Z-slides (0.5 μm step-size) of fluorescence is shown. (B) Quantitation of the T-shaped cells in the tea1Δ mutant treated with DMSO (Unt.), KCl 0.6 M, sorbitol 1.2 M, or 37°C (heat) for 1 hour, then washed and allowed to grow without the drug for 3 hours. The graph represents the mean ± SD of > 200 cells from two independent experiments. (C) Quantitation of T-shaped wild-type, tea1Δ, tea1Δ rgf1Δ, and tea1Δ rgf1ΔPTTR cells grown to log phase in YES liquid medium at 28°C. The graph represents the mean ± SD of > 200 cells from two independent experiments. (D) Cells expressing rgf1-GFP and crn1-GFP (actin patches) or mCherry-atb2 (microtubules) cultured separately; mixed; and then treated with DMSO, LatA 100 μM, or MBC 50 μM for 15 minutes. (E) LifeAct-mCherry (actin) and Rgf1-GFP localization in wild-type cells treated with KCl 0.6 M for 1 hour and then washed and allowed to grow without stress for the indicated times. The maximum-intensity projection of four Z-slides (0.6 μm step-size) of fluorescence is shown. Statistical significance was calculated using a two-tailed unpaired Student’s t test. **P < 0.01; ****P < 0.0001. Scale bar 2 μm.

**S1 Movie. Tea4 and microtubule dynamics in wild-type cells.** Tea4-GFP (green) and mCherry-Atb2 (red) localization in wild-type cells. Protein dynamics was followed for 8 minutes, with pictures taken every 20 seconds. The maximum-intensity projection of six Z-slides (0.5 µm step-size) is shown.

**S2 Movie. Tea4 and microtubule dynamics in *rgf1*Δ cells.** Tea4-GFP (green) and mCherry-Atb2 (red) localization in *rgf1*Δ cells. Protein dynamics was followed for 8 minutes, with pictures taken every 20 seconds. The maximum-intensity projection of six Z-slides (0.5 µm step-size) is shown.

**S3 Movie. Rgf1 and actin dynamics in wild-type cells during osmotic stress recovery.** Rgf1-GFP (green) and LifeAct-mCherry (actin in red) localization in wild-type cells treated with KCl 0.6 M for 1 hour and then washed and allowed to grow without stress. Protein dynamics was followed for 42 minutes, with pictures taken every 3 minutes. The maximum-intensity projection of four Z-slides (0.6 µm step-size) is shown.

**S4 Movie. Rgf1 and actin dynamics in *tea1*Δ cells during osmotic stress recovery.** Rgf1-GFP (green) and LifeAct-mCherry (actin in red) localization in *tea1*Δ cells treated with KCl 0.6 M for 1 hour and then washed and allowed to grow without stress. Protein dynamics was followed for 66 minutes, with pictures taken every 3 minutes. The maximum-intensity projection of four Z-slides (0.6 µm step-size) is shown.

## REFERENCES

1. Allam AH, Charnley M, Russell SM. Context-Specific Mechanisms of Cell Polarity Regulation. J Mol Biol. 2018 Sep;430(19).

2. Drubin DG, Nelson WJ. Origins of cell polarity. Cell. 1996;84:335–44.

3. Hoffman CS, Wood V, Fantes PA. An Ancient Yeast for Young Geneticists: A Primer on the Schizosaccharomyces pombe Model System. Genetics [Internet]. 2015 Oct;201(2):403 LP--423. Available from: http://www.genetics.org/content/201/2/403.abstract

4. Chang F, Martin SG. Shaping fission yeast with microtubules. Cold Spring Harb Perspect Biol [Internet]. 2009;1(1):a001347. Available from: https://www.ncbi.nlm.nih.gov/pubmed/20066076

5. Huisman SM, Brunner D. Cell polarity in fission yeast: a matter of confining, positioning, and switching growth zones. Semin Cell Dev Biol [Internet]. 2011;22(8):799–805. Available from: https://www.ncbi.nlm.nih.gov/pubmed/21803169

6. Martin SG. Microtubule-dependent cell morphogenesis in the fission yeast. Trends Cell Biol. 2009;19:447–54.

7. Mitchison JM, Nurse P. Growth in cell length in the fission yeast Schizosaccharomyces pombe. J Cell Sci [Internet]. 1985;75:357–76. Available from: https://www.ncbi.nlm.nih.gov/pubmed/4044680

8. Bähler J, Pringle JR. Pom1p, a fission yeast protein kinase that provides positional information for both polarized growth and cytokinesis. Genes Dev. 1998;12:1356–70.

9. Martin SG, McDonald WH, Yates 3rd JR, Chang F. Tea4p links microtubule plus ends with the formin for3p in the establishment of cell polarity. Dev Cell [Internet]. 2005;8(4):479–91. Available from: https://www.ncbi.nlm.nih.gov/pubmed/15809031

10. Mata J, Nurse P. tea1 and the microtubular cytoskeleton are important for generating global spatial order within the fission yeast cell. Cell [Internet]. 1997;89(6):939–49. Available from: https://www.ncbi.nlm.nih.gov/pubmed/9200612

11. Tatebe H, Shimada K, Uzawa S, Morigasaki S, Shiozaki K. Wsh3/Tea4 is a novel cell-end factor essential for bipolar distribution of Tea1 and protects cell polarity under environmental stress in S. pombe. Curr Biol [Internet]. 2005;15(11):1006–15. Available from: https://www.ncbi.nlm.nih.gov/pubmed/15936270

12. Browning H, Hayles J, Mata J, Aveline L, Nurse P, McIntosh JR. Tea2p is a kinesin-like protein required to generate polarized growth in fission yeast. J Cell Biol. 2000;151:15– 27.

13. Browning H, Hackney DD, Nurse P. Targeted movement of cell end factors in fission yeast. Nat Cell Biol [Internet]. 2003/08/02. 2003;5(9):812–8. Available from: http://www.ncbi.nlm.nih.gov/pubmed/12894167

14. Brunner D, Nurse P. CLIP170-like tip1p spatially organizes microtubular dynamics in fission yeast. Cell. 2000;102:695–704.

15. Busch KE, Hayles J, Nurse P, Brunner D. Tea2p kinesin is involved in spatial microtubule organization by transporting tip1p on microtubules. Dev Cell. 2004;6:831–43.

16. Snaith H, Sawin KE. Fission yeast mod5p regulates polarized growth through anchoring of tea1 at the cell tips. Nature. 2003;423:647–51.

17. Snaith HA, Samejima I, Sawin KE. Multistep and multimode cortical anchoring of tea1p at cell tips in fission yeast. EMBO J [Internet]. 2005/10/14. 2005;24(21):3690–9. Available from: http://www.ncbi.nlm.nih.gov/pubmed/16222337

18. Dodgson J, Chessel A, Yamamoto M, Vaggi F, Cox S, Rosten E, et al. Spatial segregation of polarity factors into distinct cortical clusters is required for cell polarity control. Nat Commun [Internet]. 2013/05/16. 2013;4:1834. Available from: http://www.ncbi.nlm.nih.gov/entrez/query.fcgi?cmd=Retrieve&db=PubMed&dopt=Citation&list_uids=23673619

19. Feierbach B, F. V, Chang F. Regulation of a formin complex by the microtubule plus end protein tea1p. J Cell Biol. 2004;165(697–707).

20. Bi E, Park HO. Cell polarization and cytokinesis in budding yeast. Genetics [Internet]. 2012;191(2):347–87. Available from: http://www.ncbi.nlm.nih.gov/pubmed/22701052

21. Hodge RG, Ridley AJ. Regulating Rho GTPases and their regulators. Nat Rev Mol Cell Biol [Internet]. 2016;17(8):496–510. Available from: https://www.ncbi.nlm.nih.gov/pubmed/27301673

22. Martin SG, Arkowitz RA. Cell polarization in budding and fission yeasts. FEMS Microbiol Rev [Internet]. 2014;38(2):228–53. Available from: https://www.ncbi.nlm.nih.gov/pubmed/24354645

23. Machacek M, Hodgson L, Welch C, Elliott H, Pertz O, Nalbant P, et al. Coordination of Rho GTPase activities during cell protrusion. Nature. 2009 Sep 19;461(7260).

24. Bendezu FO, Vincenzetti V, Vavylonis D, Wyss R, Vogel H, Martin SG. Spontaneous Cdc42 polarization independent of GDI-mediated extraction and actin-based trafficking. PLoS Biol [Internet]. 2015;13(4):e1002097. Available from: http://www.ncbi.nlm.nih.gov/pubmed/25837586

25. Haupt A, Ershov D, Minc N. A Positive Feedback between Growth and Polarity Provides Directional Persistency and Flexibility to the Process of Tip Growth. Current Biology. 2018 Oct;28(20).

26. Johnson JM, Jin M, Lew DJ. Symmetry breaking and the establishment of cell polarity in budding yeast. Curr Opin Genet Dev [Internet]. 2011;21(6):740–6. Available from: http://www.ncbi.nlm.nih.gov/pubmed/21955794

27. Arellano M, Duran A, Perez P. Localisation of the Schizosaccharomyces pombe rho1p GTPase and its involvement in the organisation of the actin cytoskeleton. J Cell Sci. 1997;110(20):2547–55.

28. Garcia P, Tajadura V, Garcia I, Sanchez Y. Rgf1p is a specific Rho1-GEF that coordinates cell polarization with cell wall biogenesis in fission yeast. Mol Biol Cell [Internet]. 2006;17(4):1620–31. Available from: https://www.ncbi.nlm.nih.gov/pubmed/16421249

29. Garcia P, Garcia I, Marcos F, de Garibay GR, Sanchez Y. Fission yeast rgf2p is a rho1p guanine nucleotide exchange factor required for spore wall maturation and for the maintenance of cell integrity in the absence of rgf1p. Genetics [Internet]. 2009;181(4):1321–34. Available from: https://www.ncbi.nlm.nih.gov/pubmed/19189958

30. Davidson R, Laporte D, Wu JQ. Regulation of Rho-GEF Rgf3 by the arrestin Art1 in fission yeast cytokinesis. Mol Biol Cell [Internet]. 2014/12/05. 2015;26(3):453–66. Available from: https://www.ncbi.nlm.nih.gov/pubmed/25473118

31. Morrell-Falvey JL, Ren L, Feoktistova A, Haese GD, Gould KL. Cell wall remodeling at the fission yeast cell division site requires the Rho-GEF Rgf3p. J Cell Sci [Internet]. 2005;118(Pt 23):5563–73. Available from: https://www.ncbi.nlm.nih.gov/pubmed/16291723

32. Mutoh T, Nakano K, Mabuchi I. Rho1-GEFs Rgf1 and Rgf2 are involved in formation of cell wall and septum, while Rgf3 is involved in cytokinesis in fission yeast. Genes Cells [Internet]. 2005;10(12):1189–202. Available from: https://www.ncbi.nlm.nih.gov/pubmed/16324155

33. Tajadura V, Garcia B, Garcia I, Garcia P, Sanchez Y. Schizosaccharomyces pombe Rgf3p is a specific Rho1 GEF that regulates cell wall beta-glucan biosynthesis through the GTPase Rho1p. J Cell Sci [Internet]. 2004;117(Pt 25):6163–74. Available from: https://www.ncbi.nlm.nih.gov/pubmed/15546915

34. García P, Celador R, Pérez-Parrilla J, Sánchez Y. Fission Yeast Rho1p-GEFs: From Polarity and Cell Wall Synthesis to Genome Stability. Int J Mol Sci. 2022 Nov 11;23(22).

35. Pérez P, Cortés JCG, Cansado J, Ribas JC. Fission yeast cell wall biosynthesis and cell integrity signalling. Cell Surf [Internet]. 2018 Dec 1 [cited 2021 Jan 18];4:1–9. Available from: http://www.ncbi.nlm.nih.gov/pubmed/32743131

36. Garcia P, Tajadura V, Sanchez Y. The rho1p exchange factor Rgf1p signals upstream from the Pmk1 mitogen-activated protein kinase pathway in fission yeast. Mol Biol Cell. 2009;20(2).

37. Feierbach B, Chang F. Roles of the fission yeast formin For3 in cell polarity, actin cable formation and symmetric cell division. Curr Biol. 2001;11:1656–65.

38. Glynn JM, Lustig RJ, Berlin A, Chang F. Role of bud6p and tea1p in the interaction between actin and microtubules for the establishment of cell polarity. Curr Biol. 2001;11:836–45.

39. Niccoli T, Arellano M, Nurse P. Role of Tea1p, Tea3p and Pom1p in the determination of cell ends in Schizosaccharomyces pombe. Yeast [Internet]. 2003/12/10. 2003;20(16):1349–58. Available from: http://www.ncbi.nlm.nih.gov/pubmed/14663827

40. Bicho CC, Kelly DA, Snaith HA, Goryachev AB, Sawin KE. A catalytic role for Mod5 in the formation of the Tea1 cell polarity landmark. Curr Biol. 2010;20:1752–7.

41. Bähler J, Steever AB, Wheatley S, Wang Y, Pringle JR, Gould KL, et al. Role of polo kinase and Mid1p in determining the site of cell division in fission yeast. J Cell Biol. 1998;143:1603–16.

42. Yu JW, Mendrola JM, Audhya A, Singh S, Keleti D, DeWald DB, et al. Genome-Wide Analysis of Membrane Targeting by S. cerevisiae Pleckstrin Homology Domains. Mol Cell. 2004 Mar;13(5).

43. Singh N, Reyes-Ordoñez A, Compagnone MA, Moreno JF, Leslie BJ, Ha T, et al. Redefining the specificity of phosphoinositide-binding by human PH domain-containing proteins. Nat Commun. 2021 Jul 15;12(1).

44. Audhya A, Emr SD. Stt4 PI 4-Kinase Localizes to the Plasma Membrane and Functions in the Pkc1-Mediated MAP Kinase Cascade. Dev Cell. 2002 May;2(5).

45. Snider CE, Willet AH, Chen JS, Arpağ G, Zanic M, Gould KL. Phosphoinositide-mediated ring anchoring resists perpendicular forces to promote medial cytokinesis. Journal of Cell Biology. 2017 Oct 2;216(10).

46. Sawin KE, Snaith HA. Role of microtubules and tea1p in establishment and maintenance of fission yeast cell polarity. J Cell Sci [Internet]. 2004/01/22. 2004;117(Pt 5):689–700. Available from: http://www.ncbi.nlm.nih.gov/pubmed/14734657

47. Robertson AM, Hagan IM. Stress-regulated kinase pathways in the recovery of tip growth and microtubule dynamics following osmotic stress in *S. pombe*. J Cell Sci. 2008 Dec 15;121(24).

48. Salat-Canela C, Carmona M, Martín-García R, Pérez P, Ayté J, Hidalgo E. Stress-dependent inhibition of polarized cell growth through unbalancing the GEF/GAP regulation of Cdc42. Cell Rep. 2021 Nov;37(5).

49. Coll PM, Trillo Y, Ametzazurra A, Perez P. Gef1p, a new guanine nucleotide exchange factor for Cdc42p, regulates polarity in Schizosaccharomyces pombe. Mol Biol Cell [Internet]. 2003;14(1):313–23. Available from: https://www.ncbi.nlm.nih.gov/pubmed/12529446

50. Hirota K, Tanaka K, Ohta K, Yamamoto M. Gef1p and Scd1p, the Two GDP-GTP exchange factors for Cdc42p, form a ring structure that shrinks during cytokinesis in Schizosaccharomyces pombe. Mol Biol Cell [Internet]. 2003;14(9):3617–27. Available from: https://www.ncbi.nlm.nih.gov/pubmed/12972551

51. Moreno S, Klar A, Nurse P. [56] Molecular genetic analysis of fission yeast Schizosaccharomyces pombe. In 1991. p. 795–823. Available from: https://linkinghub.elsevier.com/retrieve/pii/007668799194059L

52. Calvo IA, García P, Ayté J, Hidalgo E. The transcription factors Pap1 and Prr1 collaborate to activate antioxidant, but not drug tolerance, genes in response to H _2_O _2_. Nucleic Acids Res. 2012;40(11).

53. García P, Coll PM, del Rey F, Geli MI, Pérez P, Vázquez de Aldana CR, et al. Eng2, a new player involved in feedback loop regulation of Cdc42 activity in fission yeast. Sci Rep. 2021;11(1).

54. Fernández-Golbano IM, Idrissi FZ, Giblin JP, Grosshans BL, Robles V, Grötsch H, et al. Crosstalk between PI(4,5)P2 and CK2 Modulates Actin Polymerization during Endocytic Uptake. Dev Cell. 2014 Sep;30(6).

